# A non-enveloped arbovirus released in lysosome-derived extracellular vesicles induces super-infection exclusion

**DOI:** 10.1101/2020.06.11.146357

**Authors:** Thomas Labadie, Polly Roy

## Abstract

Recent developments on extracellular vesicles (EVs) containing multiple virus particles challenge the rigid definition of non-enveloped viruses. However, how non-enveloped viruses hijack cell machinery to promote non-lytic release in EVs, and their functional roles, remain to be clarified. Here we used Bluetongue virus (BTV) as a model of a non-enveloped arthropod-borne virus and observed that the majority of viruses are released in EVs, both *in vitro* and in the blood of infected animals. Based on the cellular proteins detected in these EVs, and use of inhibitors targeting the cellular degradation process, we demonstrated that these extracellular vesicles are derived from secretory lysosomes, in which the acidic pH is neutralized upon the infection. Moreover, we report that secreted EVs are more efficient than free-viruses for initiating infections, but that they trigger super-infection exclusion that only free-viruses can overcome.

**Author summary:** Recent discoveries of non-enveloped virus secreted in EVs opened the door to new developments in our understanding of the transmission and pathogenicity of these viruses. In particular, how these viruses hijack the host cellular secretion machinery, and the role of these EVs compared with free-virus particles remained to be explored. Here, we tackled these two aspects, by studying BTV, an emerging arthropod-borne virus causing epidemics worldwide. We showed that this virus is mainly released in EVs, *in vivo* and in the blood of infected animals, and that inhibition of the cell degradation machinery decreases the release of infectious EVs, but not free-virus particles. We found that BTV must neutralize the pH of lysosomes, which are important organelles of the cell degradation machinery, for efficient virus release in EVs. Our results highlight unique features for a virus released in EVs, explaining how BTV transits in lysosomes without being degraded. Interestingly, we observed that EVs are more infectious than free-virus particles, but only free-viruses are able to overcome the super-infection exclusion, which is a common cellular defense mechanism. In conclusion, our study stresses the dual role played by both forms, free and vesicular, in the virus life cycle.

## Introduction

Lipid envelopes surrounding viruses are likely the results of a convergent evolution, potentially acquired during the adaptation of non-enveloped virus to animals [1]. The envelope plays essential roles, such as the fusion with cellular membranes to enter in cells, or the assembly of new virus particles during budding at the plasma membrane. In contrast, non-enveloped viruses developed alternative mechanisms, such as cell entry by membrane penetration [2] and exit by cell lysis. In both cases, free virus particles remain the canonical unit of virus infection, with a single virus particle entering a cell to replicate and amplify its genome. Recent studies in the field of non-enveloped viruses have drawn attention on previously unknown cell to cell spread mechanisms, in which released virus particles are not naked. A seminal work on Hepatitis A virus revealed that virus particles are released both in enveloped and non-enveloped states, with the envelope protecting virus particles from neutralising antibodies [3]. More recently, it was reported that other non-enveloped lytic viruses, such as Coxsackievirus B [4], Hepatitis E virus [5], Poliovirus [6,7], Polyomavirus [8,9], Rotavirus and Norovirus [10], can also exit the cells as cloaked in extracellular vesicles (EV). Further, it was observed that EVs could contain multiple virus particles, allowing infection of a unique cell with many virus particles at a time [7,10]. This new transmission mechanism, reported as en block transmission, allows viral genome complementation during the infection and could be a regulator of fitness evolution for a virus population [11,12].

However, the mechanisms by which viruses hijack the host cellular secretion machinery, or the benefit of vesicular transmission are still unclear [11]. Understandings these might alter the rigid definition of non-enveloped viruses. To this end, here we investigated EV associated virus release using a model non-enveloped multi-layered capsid virus, Bluetongue virus (BTV), of the Orbivirus genus, with a segmented double stranded RNA genome. BTV is an insect-borne virus that periodically causes epidemics among ruminants worldwide, with a particularly high mortality rate in sheep. Although classified as a non-enveloped virus, BTV is able to exit cells by both lytic and non-lytic mechanisms [13–15].

Here, we report that EVs containing multiple virus particles are the main infectious unit of BTV. Our results revealed a mechanism of en block transmission in which both MVBs and the late stages of autophagy are essential to cloak virus particles in secreted lysosomes. We demonstrate that EVs are not limited to cell cultures and are found in the blood of animals, and explain how both forms of the virus (free and vesicular) contribute to its life cycle are thus necessary. Indeed, we observed that EV associated virus induces a more efficient infection and synthesis of viral protein than does infection with free virus particles, while free virus particles are essential to overcome super-infection exclusion.

## Results

### Extracellular vesicles containing multiple virus particles are the main infectious unit

BTV egresses infected cells not only by cell lysis but also by a non-lytic mechanism [13,15]. Since BTV particles were shown to be associated with MVBs components during trafficking [16] and the purified virus particles are associated with NS3 [15], a membrane protein, it is possible that multiple BTV particles are released within EVs. To determine this, we infected sheep cells (PT cell line) and Culicoides insect cells (KC cell line) at a multiplicity of infection (MOI) of 10, and cell culture supernatants were centrifuged to isolate free virus particles from large EVs (~1 μm) which sediment as pellets under the conditions used. Viral supernatants were harvested at 24 hours post-infection (hpi) as the released virus content is high at this time point yet there is almost no detectable cell death (S1A Fig). Virus titrations (Fig 1A) and viral genome quantification by quantitative PCR (Fig 1B) both revealed that the majority of infectious virus particles and viral genomes were sedimented after a 10,000xg centrifugation. Further, western blot analysis of the structural protein content, both in the pellet and the supernatant revealed that VP2, VP5 and VP7 were at higher levels in the pellet (Fig 1C), consistent with the virus titres. Moreover, specific infectivity, defined as the ratio of the number of genomes per PFU [17], was significantly lower for virus particles in the pellets than in the supernatant (Fig 1D), was significantly lower in the 10,000xg pellets when compared to the supernatant (Fig 1D), suggesting that the form of virus present in the pelleted fraction was more infectious than BTV present in the supernatant. To localise the BTV proteins within purified EVs, we used a super resolution microscopy approach that could visualise both structural protein VP7 and non-structural protein NS3 with the phosphocholine BODIPY, indicating that BTV is associated with lipid membranes in EVs (Fig 1E). Moreover, direct observations of both purified EVs secreted from infected cells (Fig 1F), and ultrathin sections of EVs embedded in agar (Fig 1G) by negative stain electron microscopy revealed the presence of large EVs (ranging from 0.5 to 1.5μm) containing multiple virus particles, corroborating our fluorescence imaging observations. Altogether, these data established that the majority of BTV virus particles are released in large EVs secreted from mammalian and insect infected cells *in vitro*. To further analyse the BTV particles found in the 10,000xg supernatant, this supernatant was centrifuged again at 100,000xg, and the infectivity was measured in both the 100,000xg pellet and its supernatant (Fig 1H). Most of the infectivity was then associated with the 100,000xg pellet, that was analysed by electron microscopy for the form of virus present (Fig 1I). We observed that the majority of virus was in the form of aggregates, made of particles embedded in a lipid membrane, or in the form of naked virus particles. The lipid membrane in the case of the 100,000xg pellet may be a transient lipid membrane acquired from single particles budding at the plasma membrane, as already described [14], so we refer to the 10,000xg supernatant as the free virus fraction in this report.

**Fig 1.**
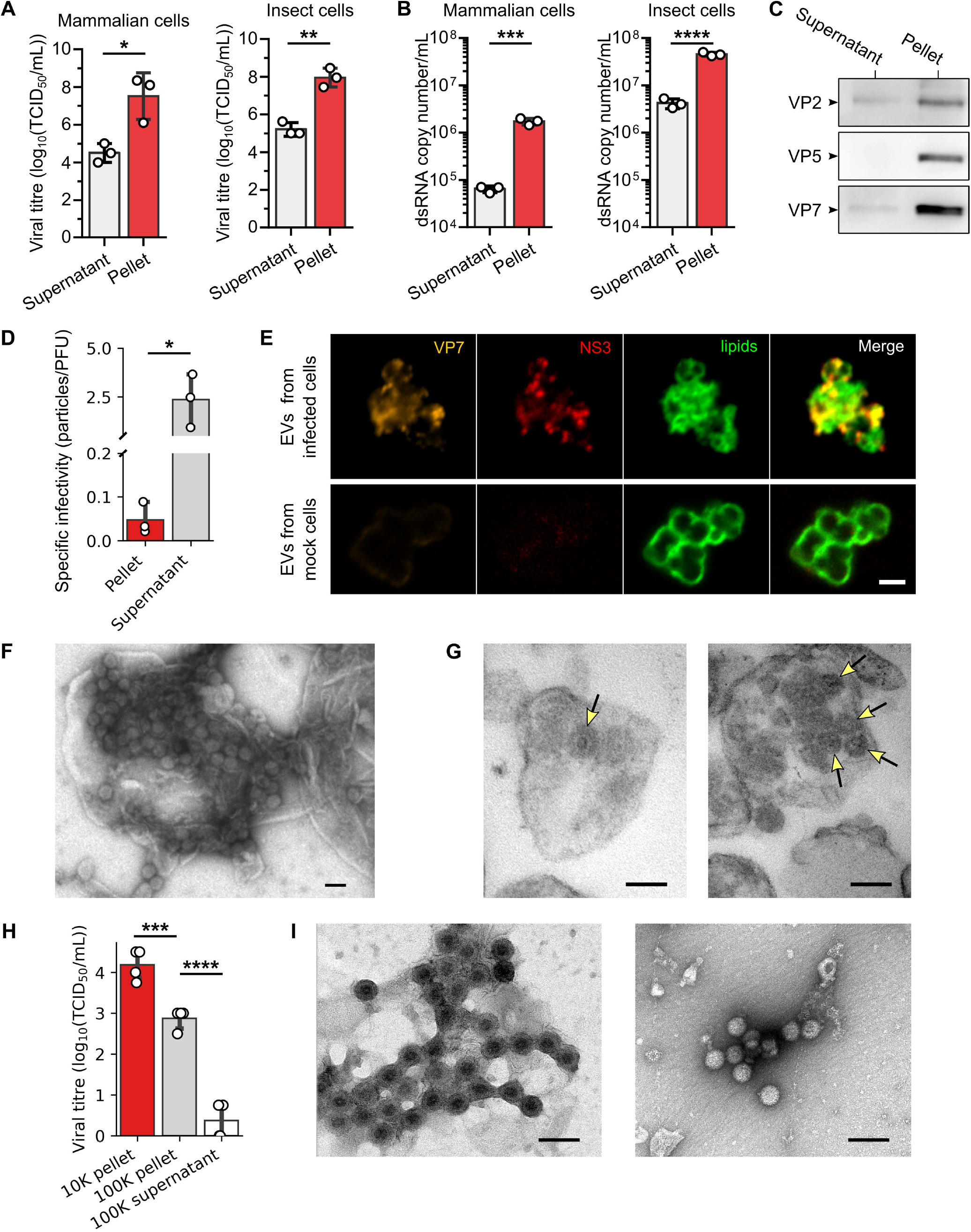
The majority of BTV virus particles are released in large EVs. (A and B) Virus particles (A) and viral genome (B) quantified in the pellet or in the supernatant after isolation of EVs secreted by infected mammalian sheep cells and insect cells. Data are presented as mean ± SD. Each point indicates the value of independent replicates (N=3; unpaired t-test, *p<0.05, **p<0.01, ***p<0.001, ****p<0.0001). (C) Outer capsid proteins VP2 and VP5, and inner capsid protein VP7, detected in the pellet and supernatant after isolation of EVs secreted by infected mammalian cells. (D) Specific infectivity, which is the number of genome copies per plaque forming unit, measured for virus particles in the pellet and the supernatant after a 10,000xg centrifugation. Data are presented as mean ± SD. Each point indicates the value of independent replicates (N=3; unpaired t-test, *p<0.05). (E) Immunofluorescence microscopy images of purified EVs secreted from infected (two top rows) or mock (bottom row) mammalian cells labelled with anti VP7 (orange) and NS3 (red) antibodies, and a fluorescent lipid membrane marker (green). Scale bar: 2 μm. (F and G) Electron microscopy images of whole EVs (F) and sectioned EVs (G) secreted from infected mammalian cells. Yellow arrows indicate the virus particles positions in the sections. Scale bar: 100nm. (H) Virus particles quantified in the pellets and the final supernatant after centrifugation of 10,000xg and 100,000xg. Data are presented as mean ± SD. Each point indicates the value of independent replicates (N=3; unpaired t-test, ****p<0.0001). (I) Representative electron microscopy images of BTV particles observed in the 100,000xg pellet. Scale bar: 100nm.

We also confirmed this phenomenon *in vivo* in bovine blood samples positive for BTV-8 (S1B Fig) collected during BTV outbreaks in France in 2019 (Fig 2A). Viral titrations of the EVs and free virus particles fractions separated by differential centrifugation revealed that virus particles were mainly contained within EVs (Fig 2B). However, no significant difference in infectivity between free virus particles and EVs was found in sheep blood samples infected with BTV-1 collected during the 2013 outbreaks. Analysis of purified bovine EVs by transmission electron microscopy showed EVs containing one or more virus particles (Fig 2C) supporting the observation that BTV virus particles are also released within EVs *in vivo*.

**Fig 2.**
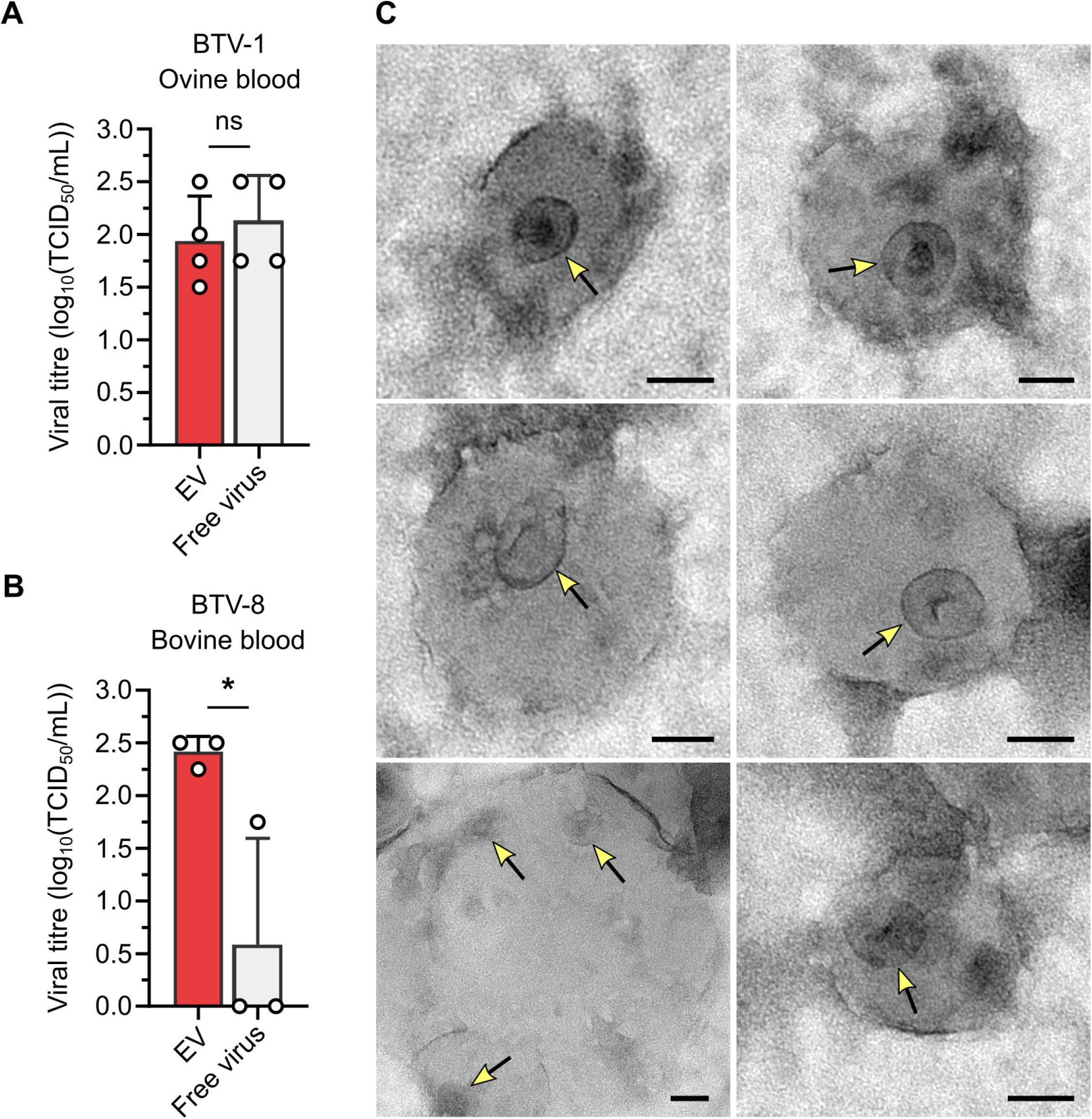
EVs containing virus particles are found in the blood of infected bovines. (A and B) Infectivity of EVs and free virus particles released from BTV-1 infected sheep blood (N=4) (A) and from BTV-8 infected bovine blood (B). Data are presented as mean ± SD. Each point indicates the value of independent replicates (N=3; unpaired t-test, ns p>0.05, *p<0.05;). (C) Electron microscopy images of EVs purified from infected bovine blood. Yellow arrows indicate virus particles. Scale bar: 50nm.

### EVs containing multiple virus particles are derived from secretory lysosomes

To determine the origin of EVs containing BTV virus particles, we tested the presence of specific cellular markers. Among the different markers of EVs assessed, we determined the presence of Annexin A2, the tumour susceptibility gene 101 (TSG101), LC3-I and LC3-II, and the lysosomal associated protein LAMP1 by western blot (Fig 3B and S2A Fig). Note that TSG101 is a MVBs marker and LC3B an autophagy related protein. To identify the cellular mechanisms involved in virus packaging in EVs, we first investigated the potential role of autophagy, using the 3-methyladenin (3-MA) that inhibits class III PI3K activity. We observed that 3-MA failed to significantly decrease the secretion of infectious EVs from sheep cells infected at a MOI of 10 (Fig 3B), suggesting that the induction of autophagy is not essential for the secretion of infectious EVs. However, inhibition of autophagosome - lysosome fusion with chloroquine (CQ), led to a significant reduction of infectious EVs released as compared to the control (Fig 3B), indicating that the late steps of autophagy are necessary for infectious EVs. In addition, inhibition of MVBs regulator protein HSP90 using geldanamycin in BTV-infected cells (MOI=10) also led to a significant reduction of infectivity measured in the EVs fraction, as compared to the control, indicating a possible role for MVBs in the release of infectious EVs. In contrast, GW4869, a drug that inhibits the release of exosomes (small vesicles ~200nm) derived from MVBs, did not affect the secretion levels of EVs containing BTV (Fig 3B). As expected, except for GW4869, these drugs also had an inhibitory effect on the virus replication (S2B Fig), affecting virus released in both EV and free viruses, albeit to various levels. These cumulative inhibitory strategies of autophagy and MVBs provide evidences that MVBs and the late stages of autophagy are both involved in the release of EVs containing virus particles. However, these EVs cannot be exosomes, based on the results obtained with the GW4869 inhibitor. Following recent description that autophagy and MVBs share common molecular machinery and organelles such as amphisomes [18], we asked if EVs could be derived from secreted lysosomes after the fusion with amphisomes. Therefore, we purified the EVs using an isopycnic ultracentrifugation and found that most of the infectivity was associated with the low-density fractions (~1.05 to 1.08 g/mL), along with the cathepsin enzymes, a lysosomal marker (Fig 3C). Interestingly, the ratio of cell surface LAMP1 to cytosolic LAMP1 was also significantly higher in infected cells (Fig 3D), indicating an increase of LAMP1 located at the plasma membrane in infected cells. The presence of viral proteins was then investigated in the lysosomes of BTV infected cells.

**Fig 3.**
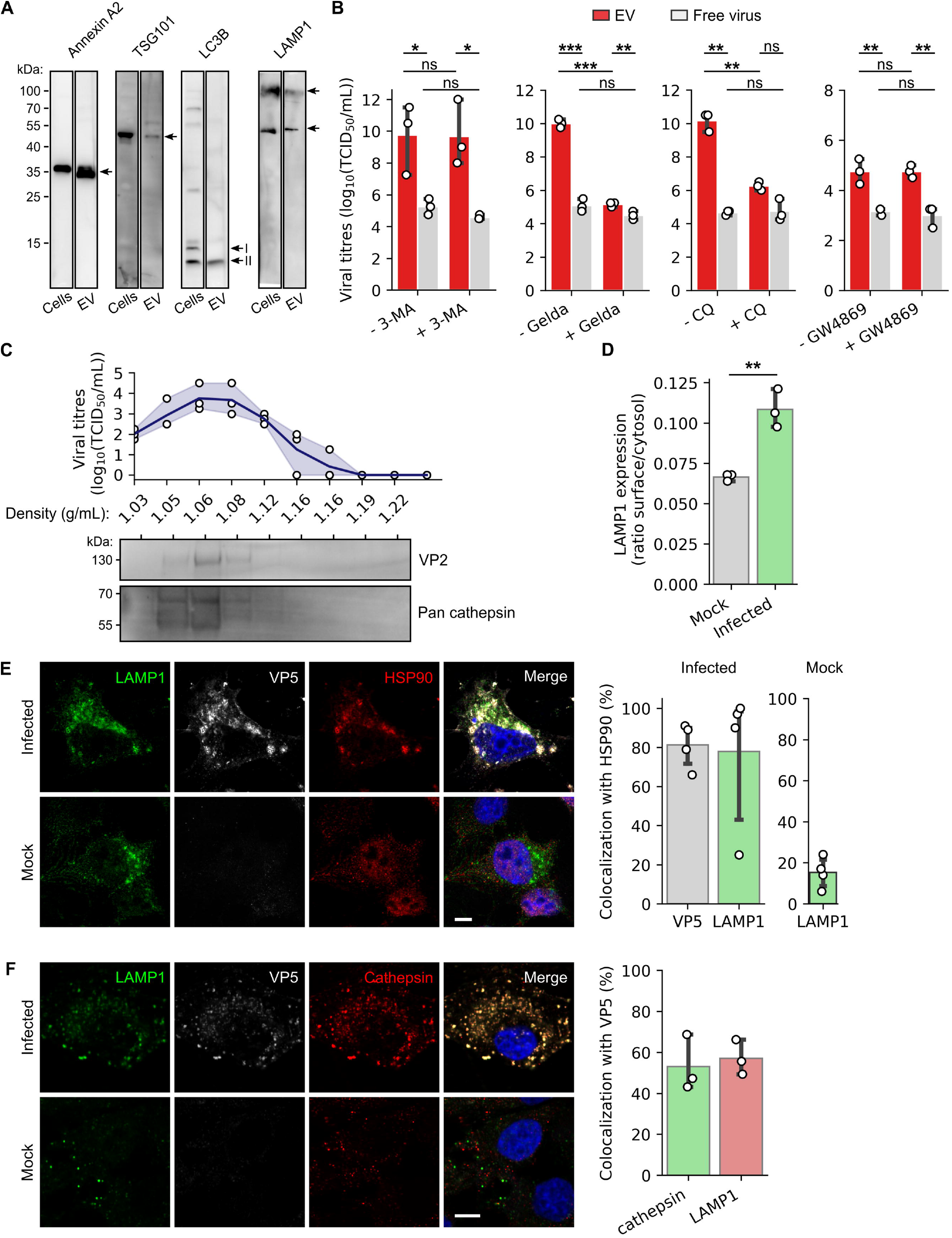
Infectious EVs rely on MVBs and late autophagy. (A) Western blot detection of cellular proteins Annexin A2, TSG101, LC3B and LAMP1 in infected sheep cell lysates and EVs secreted from these cells. (B) Infectivity of the EVs and free virus particles fractions, released from infected sheep cells in presence or absence of 3-MA, geldanamycin, CQ or GW4869. Data are presented as mean ± SD. Each point indicates the value of independent replicates (N=3, unpaired t-test, ns p>0.05, *p<0.05, **p<0.01, ***p<0.001). (C) Infectivity measured (top panel) and proteins detected (VP2 and cathepsins, bottom panel) by western blot in each fraction after isopycnic ultracentrifugation of the EVs containing BTV particles. The density (g/mL) measured in each fraction is indicated on the x-axis. Each point indicates the value of independent replicates (N=3), the continuous line represents the mean viral titre and the translucent error bands represent the confidence interval. (D) LAMP1 surface expression ratio, in mock and BTV infected cells. Data are presented as mean ± SD. Each point indicates the value of independent replicates (N=3, unpaired t-test, **p<0.01;). (E-F) Microscopy images of mock or BTV infected sheep cells labelled with anti LAMP1, VP5 and (E) HSP90 or (F) pan-cathepsin antibodies. Scale bar: 5 μm. Right panels show (E) the percentage of VP5 and LAMP1 co-localising with HSP90, and (F) the percentage of VP5 co-localising with cathepsins and LAMP1 in infected cells.

Using super resolution microscopy, we observed that in infected cells only, outer capsid protein VP5 as well as LAMP1 co-localised with the MVBs marker HSP90 (Fig 3E), indicating a fusion between MVBs and the autophagy compartments. Supporting these results, we also observed the co-localisation of LAMP1 and VP5 with TSG101, another MVBs marker (S2D Fig), and the co-localisation of VP5 with the lysosomal markers LAMP1 and cathepsins, indicating the presence of viral proteins within lysosomes (Figs 3F). Altogether, increased LAMP1 localisation at the plasma membrane, a hallmark of increased lysosomal exocytosis [19,20], and the presence of BTV proteins within lysosomes indicate that BTV infection could reprogram lysosomal degradation pathways to facilitate virus secretion.

NS3 is the only viral membrane protein and it is likely to be involved in the mechanism of EV secretion. Therefore, we tested the presence of virus in EVs (Fig 4A) following infection with available NS3 mutants [14,21,22]. Two of these mutants impair the synthesis of the two NS3 isoforms, respectively NS3 (mutant NS3_M1_) and NS3A (mutant NS3_M14_), while another with a modified late domain motif (NS3_GAAP_) cannot bind TSG101 and one, unable to bind the outer capsid protein VP2 (NS3_stop211_). Of the four mutant viruses, only BTV NS3_M14_ significantly reduced virus particle release in EVs (Fig 4A). To understand this phenotype, we analysed the localisation of the mutant virus particles in infected cells and observed that the outer capsid protein VP5, and NS3, were still co-localised with the lysosomal markers LAMP1 and cathepsin (Fig 4B, S3 Fig), suggesting that the absence of BTV NS3_M14_ in EVs was not due to a defect in virus trafficking. We then examined lysosomes in cells infected with wild type BTV (BTV_WT_) and non-infected cells expressing a LAMP1-GFP fusion protein in the presence of a fluorescent lysotracker specific to acidic organelles (Fig 4C). Confocal microscopy revealed a significant decrease of lysotracker fluorescence intensity in the lysosomes of BTV_WT_ infected cells, when compared to non-infected cells, suggesting that BTV_WT_ neutralises the acidic pH of lysosomes (Fig 4D). The number of lysosomes detected in infected cells was also significantly higher than in non-infected cells (Fig 3E and 3G). In contrast, cells infected with BTV NS3_M14_ showed comparable levels of lysotracker fluorescence to non-infected cells, suggesting that BTV NS3_M14_ was unable to counter lysosomes acidification. To provide further confirmation, cells infected with BTV NS3M_14_ and non-infected cells were examined in the presence of the lysosomal V-ATPase inhibitory drug bafilomycin A1 (Fig 3E and 3G). Interestingly, bafilomycin A1 significantly increased the levels of BTV NS3_M14_ in EVs (Fig 4H), while not affecting the intracellular replication of BTV NS3_M14_, whereas the drug had a strong inhibitory effect on the replication of BTV_WT_ (S2B Fig). Consistent with this model, we observed that the intracellular calcium levels of cells infected with BTV_WT_ were significantly higher than the calcium levels in BTV NS3_M14_ and non-infected cells (Fig 4I and 4J). Together, our data strongly support that newly synthesised BTV particles are released *via* EVs derived from secretory lysosomes after pH neutralisation.

**Fig 4.**
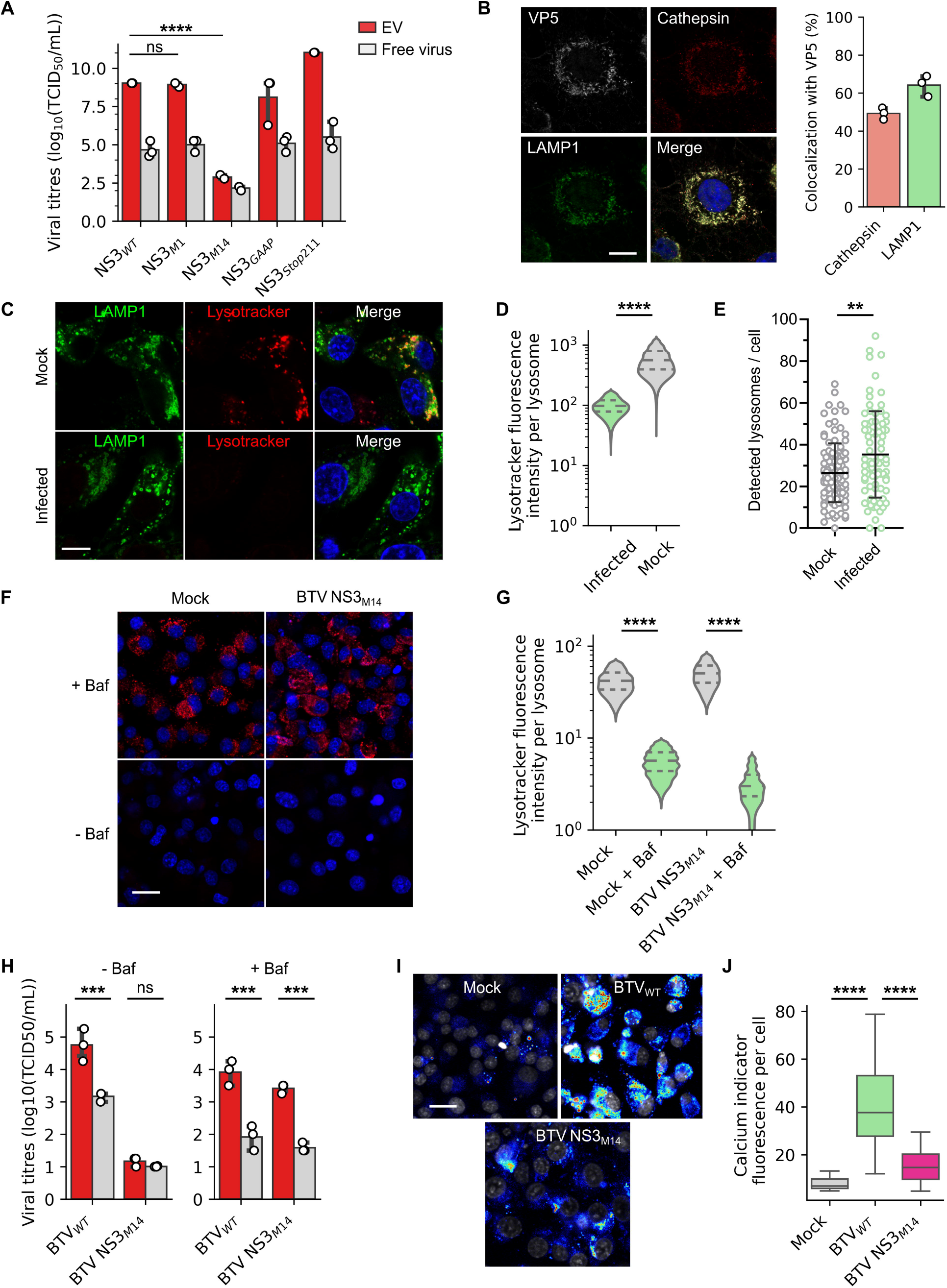
Infectious EVs are derived from lysosomal compartments. (A) Infectivity of the EVs and free virus particles fractions, released from sheep cells infected with different BTV virus with a mutation in the NS3 protein. Data are presented as mean ± SD. Each point indicates the value of independent replicates (N=3, unpaired t-test, ns p>0.05, ****p<0.0001). (B) Microscopy images of sheep cells infected with BTV NS3_M14_ and labelled with anti LAMP1, VP5 and pan-cathepsin antibodies. Scale bar: 5 μm. Right panel show the percentage of VP5 co-localising with cathepsins. (C) Microscopy images of mock and BTV_WT_ infected sheep cells (MOI =10) expressing fluorescent fusion protein LAMP1-GFP (green), in presence of an acidic compartment fluorescent marker (red). Scale bar: 5 μm. (D) Fluorescence intensity of the acidic compartment fluorescent marker detected for each lysosome, in BTV_WT_ or mock infected sheep cells. Violin plots depict min to max values of fluorescence intensity, while the horizontal lines indicate medians (solid lines) and quartiles (dashed lines) (N=4; unpaired t-test, ****p<0.0001). (E) Average numbers of lysosome/cell in BTV_WT_ infected and mock sheep cells. Data are presented as mean ± SD. Each point indicates the values of each analysed cell (N=4, unpaired t-test, **p<0.01;). (F) Microscopy images of mock and BTV NS3_M14_ infected sheep cells (MOI=10) in presence or not of bafilomycin A1, and an acidic compartment fluorescent marker (red). Scale bar: 20 μm. (G) Fluorescence intensity of the acidic compartment fluorescent marker detected for each lysosome, in BTV NS3M_14_ or mock infected sheep cells with bafilomycin A1 or not. Violin plots depict min to max values of fluorescence intensity, while the horizontal lines indicate medians (solid lines) and quartiles (dashed lines) (N=3; unpaired t-test, ****p<0.0001). (H) Infectivity of the EVs and free virus particles fractions, released from sheep cells infected with BTV_WT_ or BTV NS3_M14_ in presence (right) or absence (left) of bafilomycin A1. Data are presented as mean ± SD. Each point indicates the value of independent replicates (N=3, unpaired t-test, ns p>0.05, ***p<0.001). (I) Microscopy images of mock, BTV_WT_ and BTV NS3_M14_ infected sheep cells (MOI=10) in fluorescent calcium levels indicator (low relative fluorescence in blue and high relative fluorescence in red). Scale bar: 20 μm. (J) Fluorescence intensity of the calcium indicator in mock, BTV_WT_ and BTV NS3_M14_ infected sheep cells. Box plots depict min to max values of fluorescence intensity, while the horizontal lines indicate medians and quartiles (N=3; unpaired t-test, ****p<0.0001).

### EVs facilitate rapid infection, but only free viruses can overcome super-infection exclusion

Along with their cellular origin, the role of the secreted form of non-enveloped viruses remains unclear. To better understand the role of EVs during BTV infection, we compared the infectivity of EVs and free virus particles, by analysing the kinetic of total virus production in cells infected either by EVs or free-virus particles. Infection of mammalian cells with infectious EVs or free virus particles led to a similar amount of virus released from cells at 33 hpi. However, at 9 hpi, we found that cells infected with infectious EVs released more virus (Fig 5A), and showed a higher level of viral genome in their cytoplasm (Fig 5B) than cells infected with free virus particles. This suggests that infectious EVs are a more efficient form of infecting agents than are free virus particles. To test this possibility, we investigated the kinetic of viral protein synthesis by following the formation of viral inclusion bodies (VIBs, virus assembly factories) in the cytosol of cells infected with either EVs or free virus particles (MOI=10). We examined the size and number of VIBs in infected cells between 6 to 21 hpi, by fluorescence microscopy (Fig 5C). A frequency analysis of the VIBs surface area distribution (Fig 5D and 5E) revealed that at 6 hpi, VIBs median surface areas were 3.34±1.80μm^2^ in free-virus infected cells against 2.75±0.81μm^2^ in EVs infected cells. At 12 hpi, mean VIB size surface was clearly smaller in free-virus infected cells with 3.10±1.52μm^2^ as compared to EVs infected cells (4.00±1.86μm^2^). This result suggests that virus-directed protein synthesis is considerably more efficient in EVs infected cells. An analysis of the frequency distribution of VIBs formed per cell at 6 hpi (Fig 5F) indicated that EVs infected cells had a geometric mean of 2.7 VIBs/cell whereas in free virus infected cells, the geometric mean was 1.1 VIBs/cell, supporting the idea that infectious EVs allow the formation of multiple viral factories soon after the infection. Together, these data indicate that EV infected cells produce more VIBs, that expand faster, than VIBs produced by free-virus infected cells. Thus, in addition to containing the majority of infectious virus particles released from infected cells, EVs are also more efficient than free-virus particles to initiate an infection.

**Fig 5.**
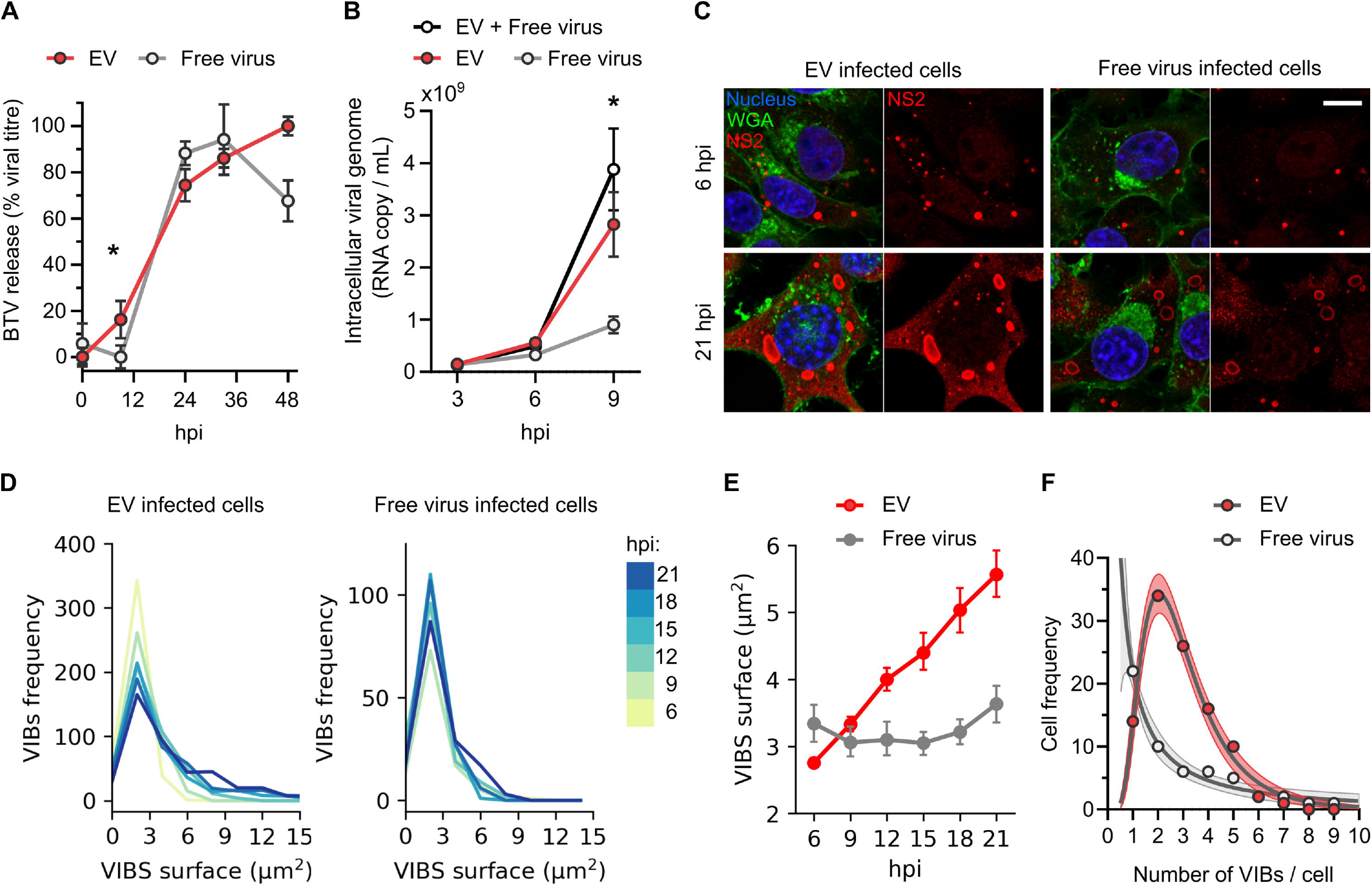
Infection with EVs is more efficient than infection with free virus particles. (A) Kinetics of virus particle release for sheep cells infected with EVs or free viruses. (B) Kinetics of intracellular viral RNA replication in sheep cells infected with EVs, free virus, or both EVs and free viruses. Each point indicates the mean relative titre (percentage of maximum reached infectivity) and vertical lines represent the SDs (N=3; unpaired t-test between EVs and free virus particles values, *p<0.05). (C) Microscopy images of sheep cells infected with infectious EVs (left panel) or free virus particles (right panel) at 6 hpi (upper panel) or 21 hpi (lower panel). For each condition a merge of the plasma membrane fluorescent marker (WGA, green) with nucleus staining (blue) and viral NS2 labelling (red), or the NS2 labelling only is shown. Scale bar: 10 μm. (D) Frequency distribution of VIB surfaces (μm^2^) measured in EVs or free virus particles infected cells between 6 and 21 hpi. (E) Kinetics of the increase in VIBs surface area in sheep cells infected with either EVs or free virus particles, derived from the frequency distribution shown in (D). Points indicate the mean surface and vertical lines represent the confidence interval. Multiple t-test showed that VIBs mean surfaces in EVs and free virus infected cells were significantly different at all time points (****p<0.0001). (F) Frequency distribution of the VIBs number/cells, after 6 hours of infection with EVs or free virus particles. Points represent the number of cells, lines represent the log normal regression analysis of the cell distributions (R^2^ = 0.98) and the shaded area represent the confidence interval (95%).

We next considered if differences between EVs and free virus particles are due to cell defence mechanisms, such as super-infection exclusion. Although not always understood, this cell defence mechanism in which infected cells become resistant to a second infection has been observed for several viruses. We first evaluated the presence of a super-infection exclusion mechanism in sheep cells infected with BTV, using two different viruses, a non-modified BTV (BTV_WT_) and a fluorescent BTV (BTV_UnaG_) encoding a fluorescent fusion protein VP6-UnaG. After controlling that BTV_UnaG_ virus particles are also mainly released in EVs (Fig 6A), we first infected sheep cells with BTV_WT_, followed by a second infection with BTV_UnaG_ immediately (0 hpi) or at 1, 2, 3, 4 or 5 hpi (Fig 6B). 12 hours after the initial infection with BTV_WT_, cells were then labelled with an anti VP6 antibody. UnaG fluorescence correspond to the detection of the BTV_UnaG_ virus only, whereas indirect fluorescence with VP6 antibody detects both viruses (Fig 6C), although less efficiently in the case of VP6-UnaG (Fig 6D). Using flow cytometry, we quantified UnaG fluorescence in cell population gated for VP6 positivity (from the antibody-based detection). These cells were either infected with BTV_WT_ (UnaG negative), BTV_UnaG_ (UnaG positive) or both viruses (UnaG positive). We used a constant MOI across all experiments, so that the number of BTV_UnaG_ single infected cells should remain constant. Therefore, variations of the UnaG fluorescence level reflect variations of co-infection with BTV_WT_ and BTV_UnaG_ and a superinfection exclusion of BTV_UnaG_. A strong decrease of the UnaG fluorescence was observed in cells with longer incubation times between the two infections (Fig 6E). This result provides evidence that BTV infection induces super-infection exclusion in sheep cells. We repeated this experiment by infecting sheep cells with BTV_WT_ (total virus particles), and then with free-virus particles or EVs of BTV_UnaG_. Sheep cells were first infected with BTV_WT_, and then BTV_UnaG_ free virus particles or infectious EVs. A super infection exclusion mechanism was observed in cells co-infected with BTV_WT_ and EVs containing BTV_UnaG_ (Fig 6F), with a significant decrease of UnaG fluorescence intensity dependent on the incubation times between the two infection. Conversely, no significant differences of UnaG fluorescence were detected in cells with secondary infection of BTV_UnaG_ free virus particles over time (Fig 6G). Altogether, these results indicate that while BTV infection led to a super-infection exclusion mechanism in sheep cells, BTV free virus particles were able to overcome this mechanism.

**Fig 6.**
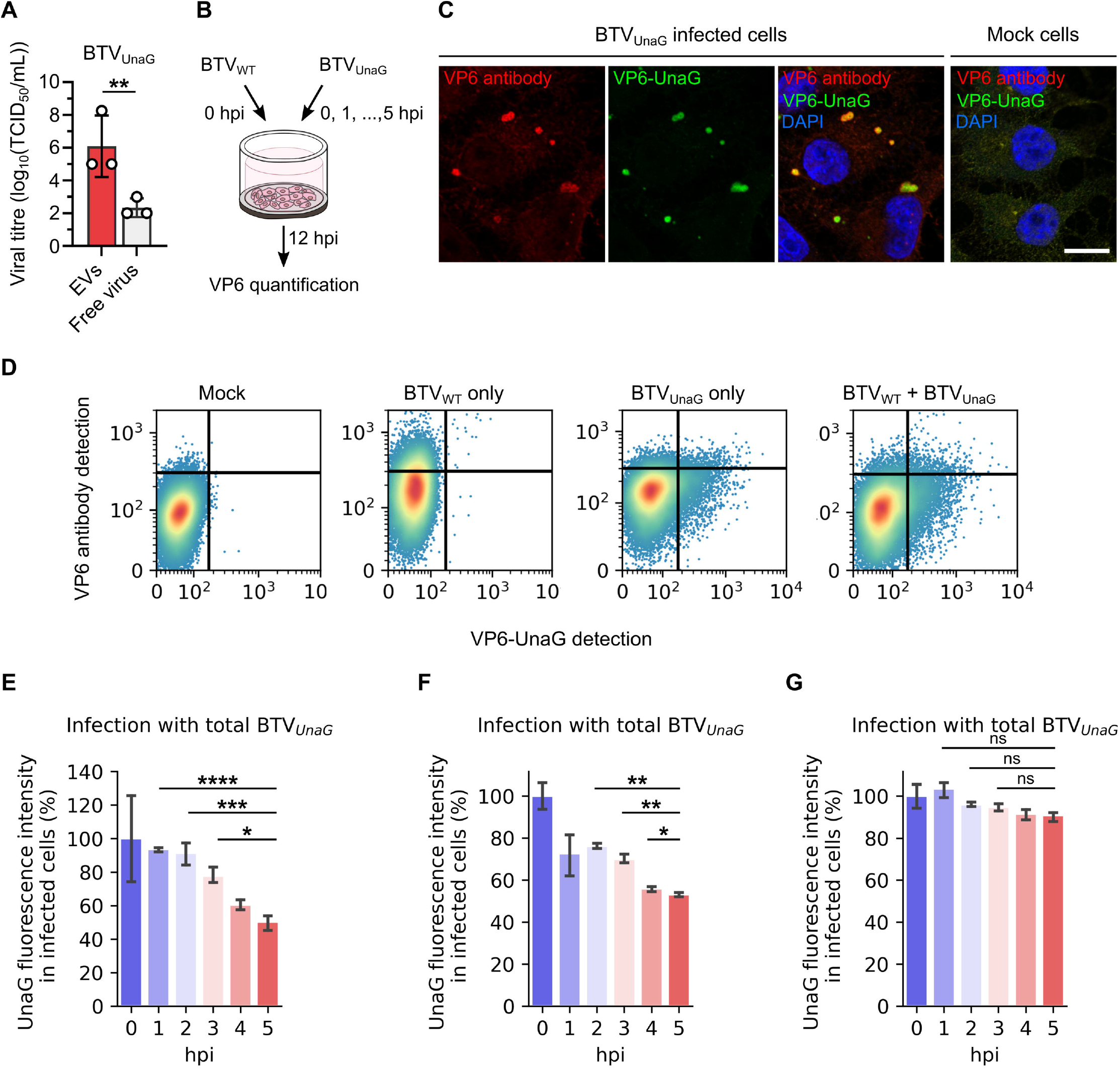
Only free virus particles overcome super-infection exclusion. (A) Infectivity of EVs and free virus particles fractions, released from sheep cells infected by BTV_UnaG_. Data are presented as mean ± SD. Each point indicates the value of independent replicates (N=3; unpaired t-test, **p<0.01). (B) Experimental design. Sheep cells were first infected with BTV_WT_, followed by BTV_UnaG_ infection at 0, 1, 2, 3, 4 or 5 hpi. VP6 quantification was performed at 12 hpi. (C) Microscopy images of mock or BTV_UnaG_ infected cells labelled with a VP6 antibody (red). VP6-UnaG (green) and nucleus (blue) were also detected. Scale bar: 10 μm. (D) Gating strategy for discrimination of non-infected sheep cells, cells infected with BTV_WT_ only, BTV_UnaG_ only, or BTV_WT_ and then BTV_UnaG_ (at 1 hpi). (E-G) Relative UnaG fluorescence (percentage of UnaG fluorescence relative to the UnaG fluorescence intensity at 0 hpi) after primary infection with BTV_WT_ and a secondary infection either with total BTV_UnaG_ (free virus particles and EVs) (E), or EVs of BTV_UnaG_ (F) or free virus particles of BTV_UnaG_ (G). Data are presented as geomean ± standard deviation factor. (N=3; ANOVA, ns p>0.05, *p<0.05, **p<0.01, ***p<0.001, ****p<0.0001).

## Discussion

While recent discoveries regarding EVs containing viruses have changed our perception of non-enveloped viruses, such observations were lacking for a non-enveloped arthropod-borne virus. Moreover, it has been hypothesised that lipid envelopes are the hallmark of virus adaptation to animals [1]. Among all the arthropod-borne viruses infecting vertebrates, arthropod-borne reoviruses are the only non-enveloped viruses. Based on these results and the recent literature on picornavirus, polyomavirus, calicivirus and rotavirus [3,7,10,23], it is tempting to suggest that the vesicular transmission could be another example of convergent evolution for a non-enveloped virus adapting to animals. In this study, we demonstrated that the majority of BTV virus particles are clustered in EVs, secreted by both mammalian and insect cells. This phenotype was observed for at least two different serotypes, BTV-1 in cell culture and BTV-8, in the blood of contaminated bovines. Despite the absence of envelope, our results highlight the presence of a lipid bilayer protecting virus particles and providing a more efficient infection initiation compared with free virus particles infection.

It has been reported that Hepatitis A virus and hepatitis E virus exit cells inside exosomes derived from the MVBs [3,5,24], that are characterised by the presence of endosomal compartments and vesicular transport markers, such as Flotillin 2, CD63 or members of the ESCRT protein family. These exosomes are around 50-200nm in size [25]. In contrast, it has been shown that Coxsackievirus B and Poliovirus are released in vesicles of approximately 500nm in diameter, through an autophagy-mediated mechanism [4,6,7]. Moreover, a recent study reported that Rotavirus egress cells in EVs derived from the plasma membrane, independently of MVBs and autophagy, and that Norovirus are release in exosomes [10]. A clear picture of different mechanisms used by non-enveloped virus for secretion in EVs is lacking. Our study contributes to understand better the diversity of release mechanisms

We showed that the lysosomal markers LAMP1 and cathepsin are present in infectious EVs, in addition to the MVB markers TSG101 and HSP90. Based on our results, it is clear that MVBs are involved in the release of BTV in EVs, because of the presence of the TSG101 in these EVs, the co-localisation of TSG101 with viral proteins in infected cells, and the deleterious effect of geldanamycin on the secretion of EVs containing virus. However, based on the size of these EVs, the failure of inhibition by GW4869, the exosome inhibitor, and the high level of infectious EVs released by the BTV NS3_GAAP_ mutant (unable to interact with TSG101 [14,22]), we conclude that the EVs release by our ovine cells cannot be exosomes. However, the presence of lysosomal markers in the EVs suggests that they could originate from the fusion between MVBs and autophagosomes that lead to the recently discovered secretory amphisome [18], virus particles could be subsequently exported in lysosomes after the amphisomes-lysosomes fusion.

Supporting this mechanism, we observed the presence of the lysosomal cathepsin proteins in the same low-density fractions as the viral particles after isopycnic density centrifugation, as well as co-localisation of the lysosomal markers LAMP1 and cathepsin with the viral proteins. Interestingly, the BTV NS3_M14_ virus, which has a mutated first start codon and is only able to synthesis an NS3A isoform of NS3 (lacking the N-terminal 13 amino-acids) [21], released significantly less virus particles in EVs than BTV_WT_. Conversely, the BTV NS3_M1_ mutant (with a mutation in the second start codon preventing the synthesis of the NS3A isoform) showed comparable levels of infectious EVs to the BTV_WT_. We did not observe a change in the co-localisation of VP5 or NS3 with the lysosomal markers in cells infected with BTV NS3_M14_ compared with BTV_WT_. In contrast, we found that lysosome acidity was higher in non-infected cells and cells infected with BTV NS3_M14_, compared with BTV_WT_. This suggests that NS3_M14_ is unable to counter lysosomal acidification as shown for WT NS3. We were able to revert this phenotype using bafilomycin A1, an inhibitor of the vacuolar type H+-ATPase, and showed that lysosomal pH disruption in BTV NS3_M14_ infected cells restored the secretion of BTV particles in EVs. This phenomenon suggests that BTV infection induces lysosomal disfunctions similar to what is observed in lysosomal storage diseases, commonly associated with an increase of lysosome biogenesis, an increased LAMP1 expression at the cell surface, as well as lysosome enlargement [26–28]. Similarly, the levels of free calcium in infected cells were lower in non-infected cells and in cells infected with BTV NS3_M14_, compared with BTV_WT_.

Interestingly, it has been proposed that the endoplasmic reticulum, in close spatial association with lysosomes, regulates the calcium storage in these organelle [29,30]. We previously showed that NS3 transits though the endoplasmic reticulum (ER), and that disrupting NS3 trafficking through the ER causes release of immature particles. It would be thus interesting to explore the interplay between NS3, the ER and lysosomes in order to determine how NS3 could regulate the secretion of BTV particles in EVs. Altogether, our data support an original model of a virus particles release mechanism involving MVBs and secretory lysosomes. Interestingly, this phenomenon is not restricted to viruses, as it has also been described for intracellular uropathogenic E. coli [20]. Related with our results, a recent preprint describes how beta-coronaviruses, which are enveloped viruses, could exploit the lysosomes for virus egress by disrupting lysosomal acidification [31]. These results, and ours, reveal a potentially unanticipated wide diversity of egress mechanism using lysosomes by different viral families.

Additionally, we also showed that EVs containing virus particles offer an efficient platform for entry and infection of naive cells. However, we observed that BTV infection triggers a superinfection exclusion phenomenon for EVs containing virus particles, whereas free virus particles can overcome it, underlying the importance of both viral forms during BTV cycle. Previously, we showed BTV VP2, the outer capsid spike binds sialic acid, a likely receptor for the free virus particles [32]. However, reports from others showed that infection with virus particles cloaked in EVs did not necessarily rely on their known virus receptor [7,8,23,33]. Here, we discovered evidences of a superinfection exclusion mechanism during BTV infectious, supporting the existence of a specific EV receptor. We observed that only free virus particles could enter in cells already infected, suggesting that EVs and free virus particles use different entry routes. In such a scenario, the receptor is not required to interact with viral outer capsid proteins, as EV secretion is already an efficient cell-to-cell communication system [34]. However, the difference in virus tropism would certainly have consequences on the virus pathogenicity. It is thus necessary to study the impact of infectious EVs on BTV induced pathogenicity, as infected ruminants variably respond to BTV infection, with cattle being mainly asymptomatic and sheep exhibiting more severe clinical signs, such as haemorrhage and ulcer in the gastro-intestinal tract or muscle necrosis [35]. In this study, we were able to identify infectious EVs in the blood of bovines infected in 2019, but it was not possible to obtain any evidence of infectious EVs in blood samples of infected sheep. However, we note that extracting EVs from frozen whole blood could potentially alter the integrity and the shape of the observed membranes structure, and further experiments will be needed to explore these aspects in more detail. It is important to note that BTV pathogenesis is markedly different in sheep and cattle and is also dependent on the virus serotype [36]. It is therefore possible that other release mechanisms dependent on the MVBs pathway, such as single virion budding at plasma membrane [16,21], are more preeminent in some hosts or with specific BTV strains. This could explain why the vesicles observed in bovine bloods tended to be smaller than the vesicles derived from the sheep cells experiments.

In natural hosts, BTV transmission requires blood feeding midges (Culicoides species) to take a blood meal from an infected animal. Our results demonstrated that Culicoides cells, for which BTV is not cytopathic, also secrets EVs containing virus particles. It would be thus interesting to investigate whether the uptake of EVs secreted by infected insects and EVs secreted by infected mammals can infect in mammals or insects respectively.

Overall, the findings described here highlight an original aspect of virus release leading to the secretion of a non-enveloped virus in host-derived secretory lysosomes, of which the acidic pH is neutralized upon the infection. Such mechanism is distinct to the proposed mechanism for the release of rotavirus in EVs [10], suggesting a divergent evolution of insect-borne double-stranded RNA viruses. Investigation of host-pathogen interactions for insect-borne viruses can significantly enhance our knowledge on the diversity of virus-containing extracellular vesicles origins. This may highlight different cellular tropisms and immunity evasion strategies between the free and vesicular forms of non-enveloped viruses and can have important implications for the development of innovative approaches for virus control.

## Acknowledgements

The authors are very grateful to Stephan Zientara and Corinne Sailleau for providing the animal blood samples (BTV national reference centre, ANSES, Maisons-Alfort, France). We also thank Eiko Matsuo (Kobe University, Kobe City, Japan) for providing the BTV_UnaG_ fluorescent virus. We acknowledge microscopy support from Mark Turmain (The Biosciences EM core facility, UCL, London UK). This research was funded by a Wellcome Trust senior investigator award to Polly Roy (100218/Z/12/Z).

## Author contribution

TL generated all the experimental data. TL and PR interpreted the data and wrote the manuscript.

## Material and Methods

### Cell lines and virus stocks

BSR cells (a derivative of the Baby Hamster Kidney cells BHK21 [37]) and PT cells (a sheep kidney cell line [38]) were maintained in Dulbecco’s Modified Eagle’s Medium (DMEM) (Sigma Aldrich), supplemented with 5% foetal calf serum (FCS) and antibiotics (100 units/mL penicillin, 100 mg/mL streptomycin, GIBCO, Life Technologies). KC cells (A Culicoides insect derived cell line) were maintained in Schneider′s Insect Medium (Sigma Aldrich), supplemented with 10% FCS and antibiotics (100 units/mL penicillin, 100 mg/mL streptomycin, GIBCO, Life Technologies). Mammalian cells were incubated at 35°C in a humidified 5% CO2 incubator and insect cells at 28°C. BTV-1 and BTVUnaG (with a segment 9 encoding a VP6-UnaG fusion protein) stocks were obtained by infecting sheep cells at a multiplicity of 10 plaque forming units per cell (pfu/cell) for 30 min under gentle agitation at room temperature (RT). The inoculum was then removed, and cells were maintained for 48 h in DMEM with 2% FCS for 48 h at 35°C. The harvested supernatants were clarified (10 min at 500xg) twice, and stored at 4°C. All the NS3 mutants were described in previous studies [14,21,22]. Stocks for these mutants were amplified as described above for BTV_WT_, and a sanger sequencing of NS3 confirmed the presence of all studied mutations.

### Virus titration and replication kinetic

Viral infectivity was estimated using tissue culture infectious dose of 50 per mL (TCID_50_/mL) titration, for which viral suspensions were diluted from 10−1 to 10−12 in D-MEM medium and inoculated to BSR cells in 96-well plates. Titres were calculated according to the Spearman-Karber method. For analysis of replication, sheep cells were inoculated with purified EVs or free virus particles in 6-well plate (MOI=1) for 1 h at RT under gentle rotation. The inoculum was then removed and replaced by D-MEM (1% FCS). Cell culture media was harvested at 9, 24, 33 and 48 hpi, clarified by centrifugation (500xg – 10 min), and infectivity was quantified by a TCID_50_ method. PT infected cells were lysed by three cycle of freeze thawing. Lysed cells were resuspended in PBS and cell debris was spun down at 500 g for 5 min. Viral RNA extraction was then performed on the harvested supernatant as indicated below.

### Viral genome quantification

Extraction of the viral RNA was performed with a Qiamp viral RNA mini kit (Qiagen), followed by a reverse transcription using a universal degenerated oligo specific to all 5′ non-coding sequences of BTV-1 segments (BTV/Uni1; 5′ GTTAAAWHDB 3′) and the GoScript Reverse Transcription (RT) System (Promega, Madison, WI, USA). The cDNAs obtained for the segment 7 were then quantified with a real time quantitative PCR (qPCR), using the 2x SYBR Green qPCR Master Mix (Bimake, Houston, TX, USA) and segment 7 specific primers (BTV1 S7/322F, 5′ GACGCCAGAGATACCTTTTAC 3′; BTV1 S7/478R, 5′ CTTGAATCATATCCGGACCAC 3′) or segment 10 specific primers (BTV1 S10/249F, 5’ CATTCGCATCGTACGCAGAA 3’; BTV1 S10/464R, 5’ GCTTAAACGCCACGCTCATA 3’). For absolute quantification of the cDNA copy numbers, a standard curve was produced using a 10-fold serial dilution of a pT7-S7 plasmid of known concentration as a template for amplification.

### Isolation of EVs by differential centrifugation

EVs were isolated from infected cells cultured in DMEM 1% FCS (mammalian cells) or Schneider′s Insect Medium (10% FCS), except for the analysis of protein markers in EVs by western blot and immunofluorescence, for which cells were cultured in 0% FCS medium in order to avoid detection of EVs present in the FCS. To identify the optimal time point for cell culture supernatant harvesting, corresponding to the highest possible viral titre with the lowest cell mortality, we used a Cell Viability Imaging Kit based on the detection of total cells number (using DAPI staining) versus cells with a compromised plasma membranes (R37609, Thermofisher Scientific).

Following harvesting of the, EVs were purified from free virus particles using a differential centrifugation protocol adapted from one used on encephalomyocarditis virus [39]. Cell debris was first removed by two steps of low speed centrifugation at 500xg for 10 min. Large EVs were pelleted by centrifuging initial volumes of 1.5 ml at 10,000xg for 30 min. Free virus particles remained in the supernatant, and pelleted EVs were resuspended in either 50 μL or 1500 μL volumes of D-MEM without FCS, depending on the concentration needed for the subsequent analysis. For all quantitative comparisons with free viruses, a correction factors were applied if the EV resuspension volumes were not equivalent to the volume prior centrifugation. When necessary, the 100,000xg centrifugation was performed for 1h at 4°C using a SW41 TI swinging bucket rotor (Beckman Coulter). The pellet was then resuspended in 50 μL or 1500 μL of D-MEM for subsequent analysis.

EVs were purified from frozen blood samples by a similar method. First, samples were quickly defrosted at 37°C, then 500 μL volumes of blood were diluted in sterile PBS (1:3) (BR0014G, Thermofisher Scientific) and centrifuged at 500xg for 10 min, followed by a 2000xg centrifugation step for 10 min in order to completely removed lysed erythrocytes. Finally, a 10,000xg centrifugation step was performed for 30 min and the pellet was resuspended in 60 μL PBS. All centrifugation steps were performed at 4°C.

### Isopycnic gradient centrifugation of EVs

EVs purification by isopycnic gradient centrifugation was performed at 150,000xg for 4 hours at 4°C, using a SW41 TI swinging bucket rotor (Beckman Coulter). EVs resuspended in 900 μL D-MEM and diluted with 100 μL iodixanol (45%) to a final 5% concentration. EVs were layered on top of a 10-45% iodixanol discontinue gradient and harvested in 9 fractions, from the bottom of the tube using a syringe. Fraction density was calculated by measuring the absorbance at 244nm after a 1:10,000 dilution and using a standard curve obtained by serial dilution of iodixanol.

### Specific infectivity

Specific infectivity of BTV in EVs and free virus fractions correspond to the number of particles required to infect a cell. We approximated the particles/mL as the genome copy number/mL measured by qPCR using S10 specific primers after extraction and reverse transcription of the viral genome. The number of pfu/mL was obtained by virus titration using a plaque assay.

### Western Blot

Proteins were detected from either resuspended EVs or cell lysates. Cell monolayers were washed twice with a PBS solution and lysed for 30 min at 4°C with a RIPA lysis buffer, 1X protease inhibitor cocktail, and 5 mM EDTA (78438, Thermofisher scientific). Cell lysates, or EVs, were diluted in 4X NuPAGE™ LDS Sample Buffer (Thermofisher scientific) to a final 1x concentration and incubated at 95°C for 5 min. Proteins were then separated on a 10% SDS gel and blotted on a PVDF membrane. When indicated, the membranes were cut horizontally for incubation with different antibodies. The anti VP2, VP5 and VP7 antibodies (made in the Pr. Roy’s laboratory [40]), anti LC3B (NB600-138455, Novus Bio), anti annexin A2 (610069, BD Bioscience), anti TSG101 (T5701, Sigma Aldrich), anti LAMP1 (ab24170, Abcam) and anti pan-cathepsin (sc-376803, Santa Cruz Biotechnology), were incubated overnight at 4°C, followed by a 2 hours incubation with respective Horseradish peroxidase-coupled secondary antibody (anti-mouse IgG ab97023, anti-guinea pig IgG ab97155 and anti-rabbit IgG ab97051, Abcam). Luminescence was then detected using SuperSignal West Pico PLUS Chemiluminescent Substrate (Thermo Fisher Scientific). The ImageJ software was used to perform linear contrast enhancement.

### Transmission electron microscopy

To image EVs without sectioning, purified EVs were resuspended in D-MEM, adsorbed onto a carbon-coated copper grid, and stained with a 1% solution of uranyl acetate. For imaging of EVs sections, purified EVs were resuspended in 30 μL D-MEM (0% FCS) and mixed with a low volume of Glutaraldehyde 25% (final concentration = 2.5%) (16220, Electron Microscopy Sciences). After 10 min of fixation at RT, EVs were briefly warmed to 37°C, and 16 μL of EVs were dropped on 4 μL of pre-melted low-melting agar (5% in water) and kept at 42°C before the addition of EVs. Samples were quickly mixed by pipetting and stored at 4°C before sectioning. After post-fixation in 1% osmium tetroxide—0.1% sodium cacodylate, EVs were dehydrated in increasing concentrations of ethanol and embedded in epoxy resin (TAAB Laboratories Equipment Ltd., Aldermaston, United Kingdom). Ultrathin sections were stained with Reynolds lead citrate and mounted onto nickel grids. Images were visualised with a transmission electron microscope (JEM-1400, JEOL Akishima, Tokyo, Japan).

### Animal blood samples

Bovine and ovine blood samples were generously provided by the French national reference centre (Maisons-Alfort, ANSES). Sheep showing clinical signs in Corsica were sampled in 2013 [41], and bovines showing clinical signs in the south-west of France were sampled in 2019 [42]. A PCR-based diagnostic performed at the time of the blood sampling confirmed the infection of sheep with BTV-1 and bovines with BTV-8. Whole blood samples were then kept frozen until reception by us, for subsequent EVs isolation.

### Drugs-mediated inhibition of EVs release

Sheep cells in 6 well-plates were infected (MOI=10) with BTV-1, 24 hours post seeding. Viral or mock inoculum were incubated 30 min at RT under gentle rotation. The inoculum was then removed and replaced by D-MEM (1% FCS), and cells were incubated at 35°C in a humidified incubator (5% CO_2_). At 1 hpi, cell culture medium was replaced with D-MEM containing 10 mM 3-methyladenine (M9281, Sigma Aldrich), 300nM geldanamycin (InvivoGen), 20μM chloroquine (C6628, Sigma Aldrich), 5μM GW4869 (D1692, Sigma Aldrich), 1μM Bafilomycin A1 (SML1661, Sigma Aldrich), so that drugs did not interfere with viral entry. These concentrations were selected because they were already validated by previous studies on BTV from us and others [43,44]. At 24 hpi, cell supernatants were harvested for isolation of EVs and analysis of infectivity. For intracellular virus particle quantification in presence or absence of the different drugs, the media culture of PT infected cells was removed at 24 hpi, and cell monolayers were rinsed twice with PBS. Cells were then lysed by 3 successive cycles of freeze/thaw at −80°C/35°C. Cell lysates were then resuspended in 500 μL DMEM and virus particles were quantified using a TCID_50_ method.

### Quantification of cell surface proteins

Cell surface protein expression of LAMP1 was measured by immunofluorescence using flow cytometry. PT cells were seeded in 6-well plates for 24 h before infection with BTV-1 (MOI=10). After 24 hpi, cell monolayers were washed twice with PBS, re-suspended using trypsin-EDTA (Thermo Fisher Scientific). Cells used for analysing the LAMP1 cytosolic content were fixed with paraformaldehyde 2% (w/v) for 10 min and permeabilised with a PBS-Triton x100 (0.05% v/v) solution for 5 min. The cells were then blocked using bovine serum albumin (PBS-BSA 1%) for 30 min and incubated with rabbit anti LAMP1 (Ab208943, Abcam) targeting the luminal domain, or an isotope control antibody (ab172730, Abcam) for 2 hours at room temperature. Cells used for analysing the cell surface LAMP1 content were labelled with the anti LAMP1 antibody for 2 hours at 4°C prior fixation. Then all labelled cells were incubated for 2 h at RT in the presence of a species-specific A488 conjugated secondary antibodies (A21071, Thermo Fisher Scientific). After centrifugation and washing, cells were re-suspended in PBS and analysed with a BD LSR II flow cytometer (BD Biosciences, San Jose, CA, USA). For each replicate, 30,000 cells were analysed using the same parameters. Flow cytometry data were analysed with FlowJo (FlowJo LLC, Ashland, OR, USA) software. For the analysis, a gating strategy discriminating doublet cells by plotting FSC-A vs. FSC-H was used, and the threshold of background fluorescence was determined using cells labelled with the isotope control antibody (S2C Fig).

### Fluorescence microscopy

For imaging, EVs were resuspended in D-MEM (0 % FCS) + fluorescent Bodipy Phosphocholine analogue at 2 μg/mL (D3793, Thermofisher Scientific) and incubated overnight at 4°C. EVs were then dropped on poly-L-Lysine (P4707, Sigma Aldrich) pre-coated chambered #1.0 Borosilicate coverslips (Nunc Lab-Tek 155411, Thermofisher Scientific), and adsorbed for 1 hour at 35°C. The residual medium was removed, and attached EVs were fixed with PBS-PFA (2%) for 10 min at RT, rinsed with PBS and permeabilised with PBS-triton X100 (0.05%) for 5 min at RT. EVs were then rinsed twice with PBS, incubated for 1 hour at RT with a solution of PBS-BSA (1%), and incubated overnight at 4°C with primary antibodies anti VP7 and NS3 (made in Pr. Roy’s laboratory), followed by fluorophore A633 and A546-conjugated antibodies (A21071 and A11074, Thermofisher Scientific) for 2 hours at RT and mounting with aqueous mounting medium (F4680, Sigma Aldrich). For sheep cells imaging, cells were cultured in chambered tissue culture treated polymer coverslips (μ-slide Angiogenesis 81506 or μ-slide 4 well 80427, Ibidi), infected with BTV-1 (MOI=10) 24 h post-seeding, and at 1 hpi, inoculated with a baculovirus encoding LAMP1-GFP at concentration of 10 particles / cell (CellLight Lysosomes-RFP, BacMam 2.0, Thermofisher scientific). At 18 hpi (unless specified), cell monolayers were rinsed twice in PBS, fixed in PBS-PFA (2%) for 10 min, permeabilised in PBS-triton X100 (0.05%) for 5 min, incubated in PBS-BSA (1%) for 30 min. Cells were labelled with primary antibodies anti HSP90 (11405-1-AP, Proteintech), anti pan-cathepsin (sc-376803, Santa Cruz Biotechnology), anti VP5 and anti NS3 overnight at 4°C, and labelled with Alexa fluor A488, A546 and A633-conjugated antibodies (A11001, A11074 and A21071, Thermofisher scientific) for 2 hours at RT. Nuclei were labelled with a PBS-Hoescht solution at 2 μg/mL for 5 min at RT and samples were finally mounted in mounting medium (F4680, Sigma Aldrich). Labelled cells and EVs were analysed with an inverted LSM 880 confocal microscope with airyscan (Carl Zeiss Ltd.), or with an inverted widefield Nikon Ti Eclipse microscope (Nikon) for the VIBs surface analysis. For the co-localisation analysis, three independent replicates (with more than 90 cells imaged for each replicate) were analysed with a Statistical Object Distance Analysis (SODA, Icy software) on the whole cells [45].

### Lysosomal pH investigation

After infection with BTV-1 (MOI=10), cells were incubated at RT under gentle rotation for 30 min, before the inoculum was replaced with D-MEM containing baculovirus expressing LAMP1-GFP at concentration of 10 particles / cell (CellLight Lysosomes-RFP, BacMam 2.0, Thermofisher scientific). At 18 hpi, cells were incubated with Hoescht at 2 μg/mL for 5 min at RT, followed by an acidic compartment marker 100nM (Lysotracker DND199, Thermofisher scientific) for 1 min at RT. Cells were then washed twice with a PBS solution and analysed with a LSM880 confocal microscope. Images were analysed with Icy (https://icy.bioimageanalysis.org). Cells were counted by detection of stained nuclei and lysosomes by detection of the fusion LAMP1-GFP fluorescent protein. Each lysosome detected was considered as a region of interest, and Lysotracker fluorescence intensity detected in these regions were then quantified.

### Cytosolic calcium assay

After infection with BTV (MOI=10), cells were incubated at RT under gentle rotation for 30 min, before the inoculum was replaced with D-MEM. At 18 hpi, cells were incubated with Hoescht at 2 μg/mL for 5 min at RT, followed by incubation with the fluorescent labelling reagent Fluo-4 AM (at 1μM in D-MEM) for the detection of intracellular calcium (GR3332801, Abcam) for 1 hour at 35°C. Fluo-4 AM/D-MEM solution was then replaced by D-MEM only to remove any non-specific binding, and left a further 30 min at 35°C, followed by two rinsing with a PBS solution and imaged with a LSM880 confocal microscope (Fluo-4 AM excitation at 488nm). For image analysis, all cells were detected using the HK mean plug-in in Icy (https://icy.bioimageanalysis.org) applied on the Fluo-4 channel, and detected cells were considered as regions of interest in which the Fluo-4AM intensity was then measured.

### Monitoring of VIBs during the infection

Cells were infected and fixed at different times post infection as above, and labelled with an anti NS2 (made in Pr. Roy laboratory) antibody, a secondary Alexa fluor A546-conjugated antibody (A11074, Thermofisher scientific), and A488-conjugated wheat germ agglutinin as a membrane label (W11261, Thermofisher scientific) for 10 min at RT prior cells permeabilisation. Microscopy images were analysed with Icy, and VIBs were detected using the spot detector plugin, which provided NS2 spot surfaces. Cell counting was performed based on the number of nuclei detected in images. For data analysis, a cut-off was applied to keep all detected NS2 spots with a surface above 22μm^2^. The frequency distribution of VIBs surface values were performed with a bin width of 15μm^2^ and the lognormal regression of the frequency distribution of the number of VIBs per cells were performed after confirming that data were following a lognormal distribution. The total number of VIBs measured at each time points is indicated in the S1 table.

### Super-infection exclusion analysis

For superinfection exclusion analysis, sheep cells in 6 well-plates were infected with BTV-1 (BTV_WT_, MOI=5) as described above. Simultaneously (0 hpi), or at 1, 2, 3, 4 or 5 hpi, cells were inoculated with purified EVs or free virus particles of BTV_UnaG_ (MOI=5) and placed at 35°C in a humidified incubator (5% CO_2_) for 1 hour. BTV_UnaG_ inoculum was then removed and replaced with D-MEM (1% FCS). At 12 hpi (7 hpi of BTV_UnaG_ infection), cell monolayers were washed twice with PBS, re-suspended using trypsin-EDTA (Thermo Fisher Scientific), fixed with paraformaldehyde 2% (w/v) for 10 min, permeabilised with a solution of PBS-Triton x100 (0.05% v/v) for 5 min, and blocked with a PBS-BSA (1%) solution for 30 min. The cells were then incubated with an anti VP6 antibody (made in Pr. Roy’s laboratory) overnight at 4°C, followed by an incubation of 2 h at RT in the presence of a species-specific Alexa Fluor A633 conjugated secondary antibodies (A21071, Thermo Fisher Scientific). After centrifugation and washings, cells were re-suspended in PBS and analysed with a BD LSR II flow cytometer (BD Biosciences, San Jose, CA, USA). For each replicate, 30,000 cells were analysed using the same parameters. The detection of Alexa Fluor 633 fluorescence corresponded to the total VP6 detected in infected cells, and the UnaG detected fluorescence to VP6 expressed by BTV_UnaG_ infection only.

### Statistical analysis

Numerical data were analysed with Python (v 3.7.1), the Scipy library (v 1.4.1) and Prism software version 8.1.2 (Graph Pad Software, San Diego, CA, USA). For all sets of data, two-tailed t-tests or ANOVA tests were performed for statistical comparison. When necessary, outlier values removal was performed using the EllipticEnvelope from the scikit-learn library (0.23.1). Figures were prepared using Jupyter notebooks (Project Jupyter, https://jupyter.org) and Inkscape (https://gitlab.com/inkscape/inkscape)

### Data availability

All relevant data are within the manuscript and its Supporting Information files.

## Supporting information

**S1 Fig. Selection of the optimal time point for EVs harvesting from infected sheep cells and confirmation of BTV presence in blood samples**. (A) The number of dead cells versus the total number of cells was measured (left panel) in our infected cell cultures, to calculate the percentage of cell mortality (middle panel) at the time of infection (0 hpi), at 24 hpi and at 48 hpi. In parallel, we measured BTV viral titres in the pellet or the supernatant after a 10,000xg centrifugation in viral suspension harvested from infected cells at 24 hpi or 48 hpi (right panel). The 24 hpi time point represented the optimal balance between cell mortality and viral titres, and was used for all the subsequent experiments. Data are presented as mean ± SD. Each point indicates the value of independent replicates (N=3; unpaired t-test, *p<0.05). (B) The presence of BTV in was confirmed in three independent blood samples by RT-qPCR targeting the segment 7. Data are presented as mean ± SD. Each point indicates the value of independent replicates (N=3).

**S2 Fig. Effect of drugs on the viral replication and co-localisation between VP5, lysosomal and MVBs markers.** (A) Western blot detection of cellular proteins Annexin A2, TSG101, LC3B and LAMP1 in EVs released from non-infected sheep cells. (B) Intracellular viral titres measured at 24 hpi in sheep infected cells in presence or absence of 3-MA, CQ, geldanamycin, GW4869 on bafilomycin A1. Data are presented as mean ± SD. Each point indicates the value of independent replicates (N=3, unpaired t-test, ns p>0.05, **p<0.01, ****p<0.0001). (C) Determination of the background fluorescence in mock and BTV infected sheep cells using an isotope control antibody. For reference, the level of fluorescence is also represented in mock cells labelled with a LAMP1 antibody. (D) Microscopy images of mock or BTV_WT_ infected sheep cells labelled with anti LAMP1, VP5 and TSG101 antibodies. Scale bar: 5 μm. Right panel show the percentage of VP5 and LAMP1 co-localising with TSG101 in infected cells.

**S3 Fig. Co-localisation between NS3 and LAMP1 in BTV_WT_ and BTV NS3_M14_ infected cells.** Microscopy images of mock, BTV_WT_, or BTV NS3_M14_ infected sheep cells labelled with anti LAMP1 and NS3 antibodies. Scale bar: 5 μm. Right panel show the percentage of NS3_WT_ and NS3_M14_ co-localising with LAMP1 in infected cells.

**S1 Table.** Total number of lysosomes analysed per time point in sheep cells infected with EVs or free virus particles.

