## Supplementary figures and images for "A non-enveloped arbovirus released in lysosome-derived extracellular vesicles induces super-infection exclusion"

### S1 Figure

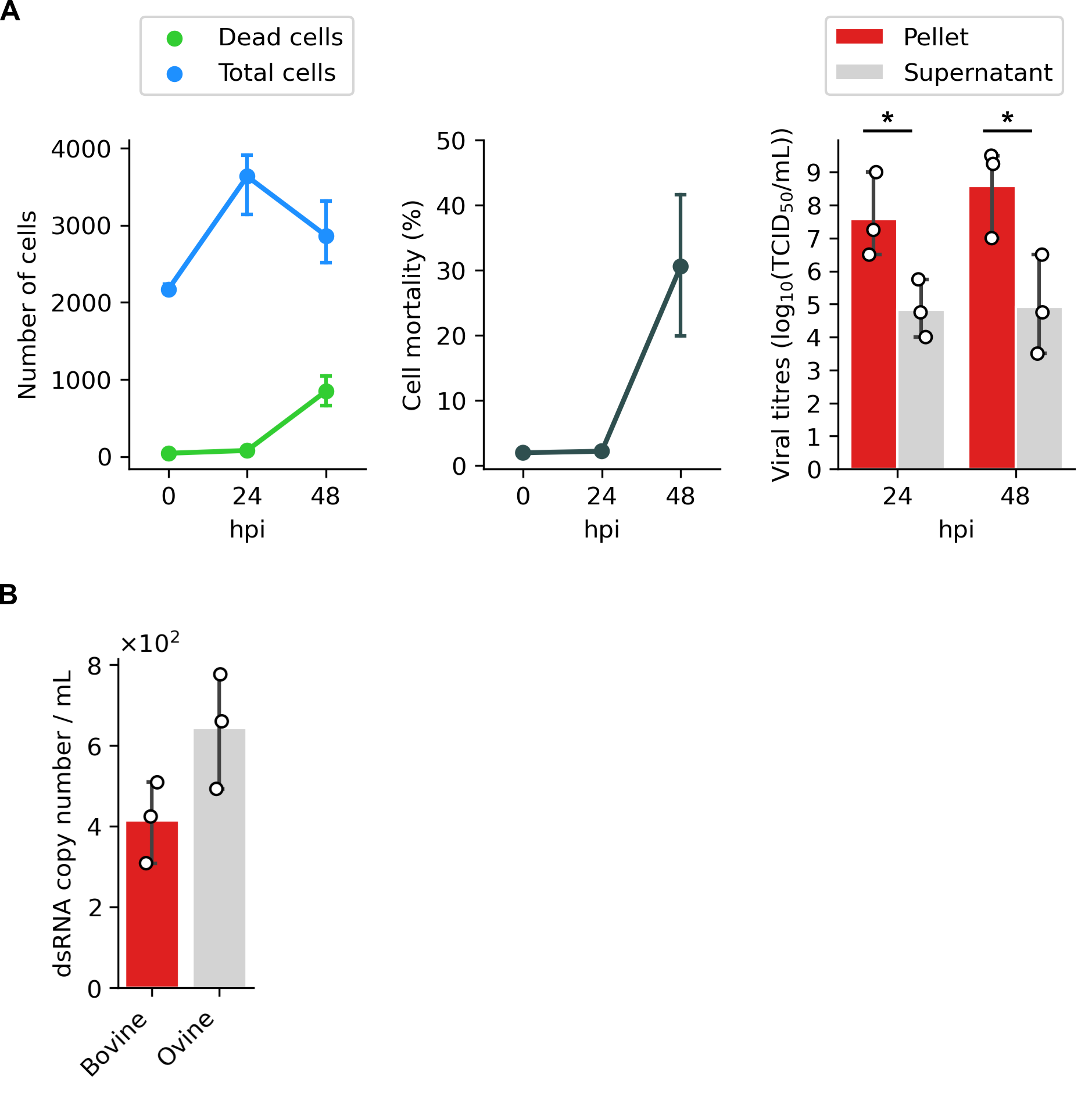

### S2 Figure

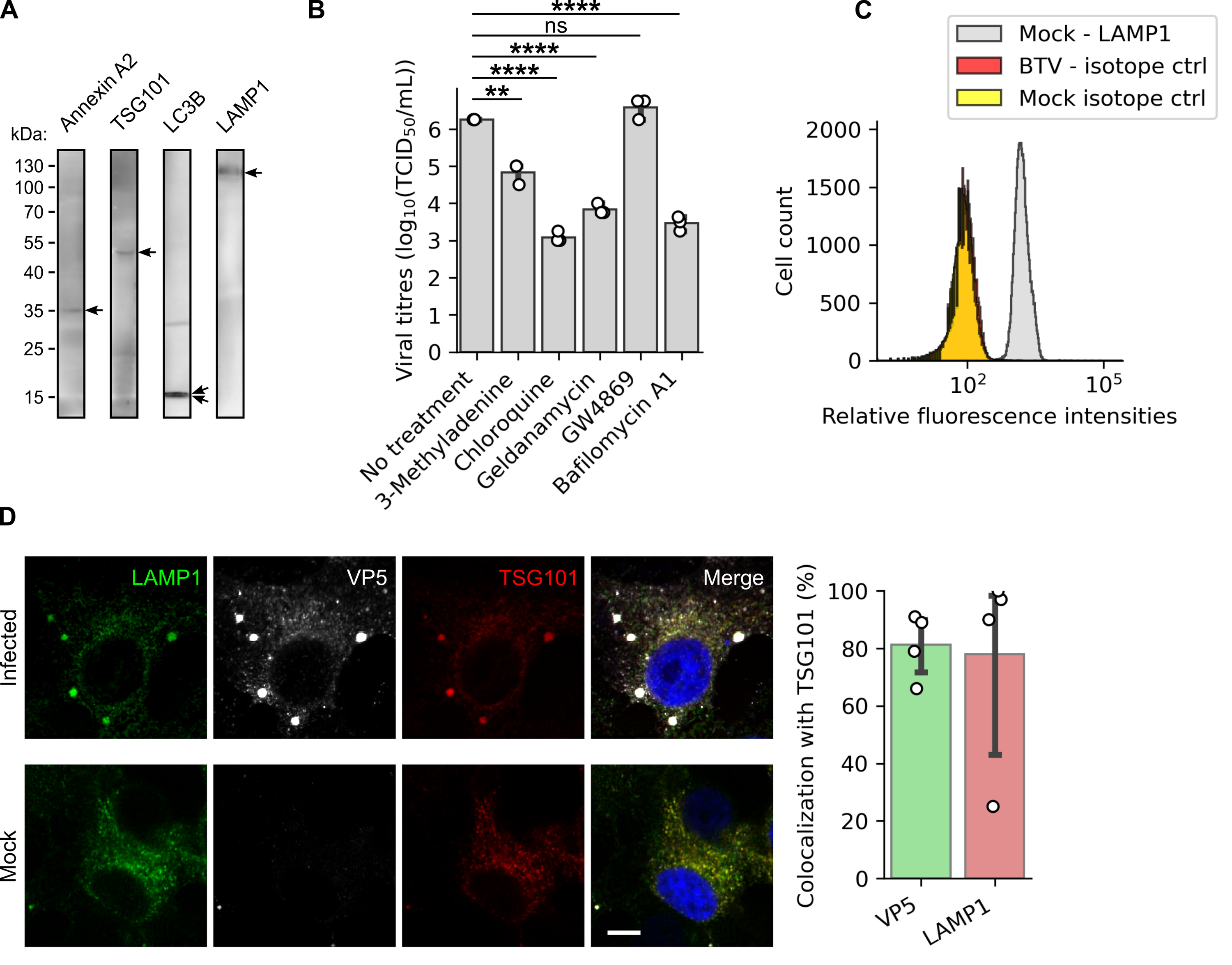

### S3 Figure

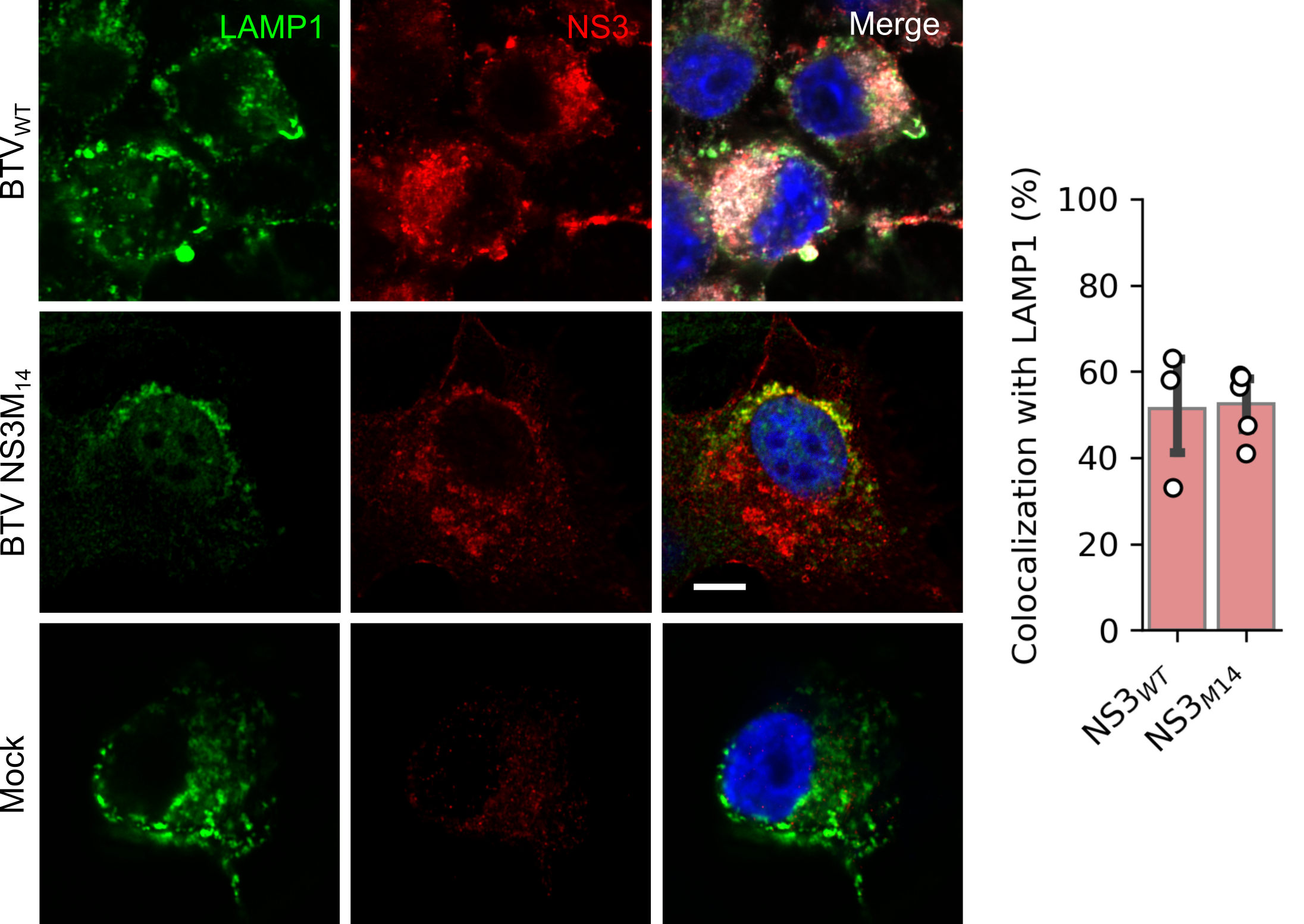
