## Supplementary material for "A non-enveloped arbovirus released in lysosome-derived extracellular vesicles induces super-infection exclusion": S1 Table

| Hpi: | 6h | 9h | 12h | 15h | 18h | 21h |
|---|---|---|---|---|---|---|
| EVs infected cells ($N_{VIBs}$) | 429 | 423 | 416 | 1020 | 982 | 912 |
| Free virus infected cells ($N_{VIBs}$) | 149 | 106 | 187 | 314 | 367 | 509 |

**S1 Table. Total number of lysosomes analysed per time point in sheep cells infected with EVs or free virus particles.**
